# Minimal impact of proprioceptive loss on implicit sensorimotor adaptation and perceived movement outcome

**DOI:** 10.1101/2023.01.19.524726

**Authors:** Jonathan S. Tsay, Anisha M. Chandy, Romeo Chua, R. Chris Miall, Jonathan Cole, Alessandro Farnè, Richard B. Ivry, Fabrice R. Sarlegna

**Author notes:** **Corresponding author Information:** Jonathan Tsay. **Funding:** JST is funded by the NIH (F31NS120448). RC is funded by the Natural Sciences and Engineering Research Council (NSERC) of Canada (2019-04513). AF is funded by ANR grants (Blind-Touch ANR-19-CE37-0005, Dec-Space ANR-21-CE28-0001 and Bodys ANR-22-CE28-0018). RBI is funded by the NIH (grants NS116883 and NS105839). FS is funded by the Mourou-Strickland mobility program (French Embassy in Canada), the NeuroMarseille Institute (French government under the Programme « Investissements d’Avenir », Initiative d’Excellence d’Aix-Marseille Université via A*Midex funding (AMX-19-IET-004), and ANR (ANR-17-EURE-0029), and the ANR grants ANR-21-CE28-0022 and ANR-22-CE33-0009.

## Abstract

Implicit sensorimotor adaptation keeps our movements well-calibrated amid changes in the body and environment. We have recently postulated that implicit adaptation is driven by a perceptual error: the difference between the desired and perceived movement outcome. According to this perceptual re-alignment model, implicit adaptation ceases when the perceived movement outcome – a multimodal percept determined by a prior belief conveying the intended action, the motor command, and feedback from proprioception and vision – is aligned with the desired movement outcome. Here, we examined the role of proprioception in implicit motor adaptation and perceived movement outcome by examining individuals who lack proprioception. We used a modified visuomotor rotation task designed to isolate implicit adaptation and probe perceived outcome throughout the experiment. Surprisingly, implicit adaptation and perceived outcome were minimally impacted by deafferentation, posing a challenge to the perceptual re-alignment model of implicit adaptation.

## Introduction

Multiple learning processes operate to ensure that motor performance remains successful in the face of changes in the environment and body. For example, if a tennis ball is consistently perturbed by the wind, the player can explicitly and rapidly adjust their swing to compensate. This perturbation will also engage an automatic, implicit adaptation process that uses the error information to recalibrate the sensorimotor system.

We have recently postulated that implicit adaptation is driven by a perceptual error, the difference between the desired and perceived movement outcome (Tsay et al., 2022) (also see: (Zhang et al., 2023)). According to this perceptual re-alignment model^*^, the perceived movement outcome is a multimodal percept determined by a prior belief conveying the intended action, the motor command, and feedback from proprioception and vision (Bhanpuri et al., 2013; Ernst & Di Luca, 2011; Sober & Sabes, 2003). In an upper limb reaching task, introducing a visual perturbation will shift the perceived outcome toward the visual cursor and, thus, away from the actual hand position and away from the target. This perceptual error would drive movements of the hand in the opposite direction to the visual perturbation (implicit adaptation). When the perceptual error is nullified, that is, when the perceived outcome is aligned with the desired outcome, implicit adaptation will cease.

Individuals lacking proprioceptive and tactile inputs from the upper limb provide an interesting test case for understanding the role of proprioception in implicit adaptation. ‘Deafferentation’ is a rare condition that arises from either a congenital disorder or a neurological insult (Bernier et al., 2006; Chesler et al., 2016; Cole & Sedgwick, 1992; Miall et al., 2018; Miller et al., 2019; Rothwell et al., 1982; Sarlegna & Sainburg, 2009; Sterman et al., 1980). Previous case studies have observed preserved motor adaptation in deafferented adults (Bernier et al., 2006; Ingram et al., 2000; Lefumat et al., 2016; Miall et al., 2018; Sarlegna et al., 2010; Yousif et al., 2015). However, the tasks used in these studies have not isolated implicit adaptation. Thus, performance changes might result from explicit, strategic processes (Ingram et al., 2000; Tsay et al., 2023). Moreover, the impact of proprioceptive loss on the perceived movement outcome during implicit adaptation is unknown.

Here, we tested a cohort of deafferented individuals on a clamped visuomotor rotation task that isolates implicit adaptation and probes perceived outcome (Morehead et al., 2017; Tsay et al., 2020). Based on the perceptual re-alignment model, we tested two core predictions. First, there should be a heightened, perceptual shift in the deafferented group compared to that of the control group. With the loss of proprioception, we predicted the deafferented group would rely heavily on vision to determine perceived outcome and thus show a heightened perceptual shift toward vision. By the perceptual re-alignment model, an increase in the perceptual shift in the deafferented group would result in heightened implicit adaptation; they will require a larger change in reach angle to offset the larger perceptual error. We test these two predictions in the following experiment.

## Results

### Implicit adaptation is preserved but not heightened in deafferented individuals

We compared the performance of six participants with a severe proprioceptive loss to that of 60 age-, gender and laterality-matched controls (10 controls matched to each deafferented participant) on a clamped visuomotor rotation task. Our task differed from prior studies of adaptation in this population in two notable ways. First, since deafferentation is less complete in the proximal muscles for a subset of the participants, we used a “reaching” task in which movement was mostly limited to wrist and/or fingers movement over a laptop computer trackpad. Second, we used clamped visual feedback, a method that isolates implicit adaptation (Morehead et al., 2017) (Figure 1A). In this task, participants reach to a visual target and receive visual cursor feedback that follows a fixed trajectory defined relative to the target. Thus, unlike standard perturbation methods, the angular position of the feedback is not contingent on the participant’s movement direction. Participants are fully informed of this manipulation and instructed to always reach directly to the target while ignoring the visual feedback. Despite these instructions, the visual perturbation between the position of the target and the cursor elicits an implicit adaptive response in healthy participants, causing a trial-by-trial change in movement direction away from the target and in the opposite direction to the cursor. These motor adjustments are not the result of explicit re-aiming; indeed, participants are oblivious to the change in their behavior (Tsay et al., 2020).

**Figure 1:**
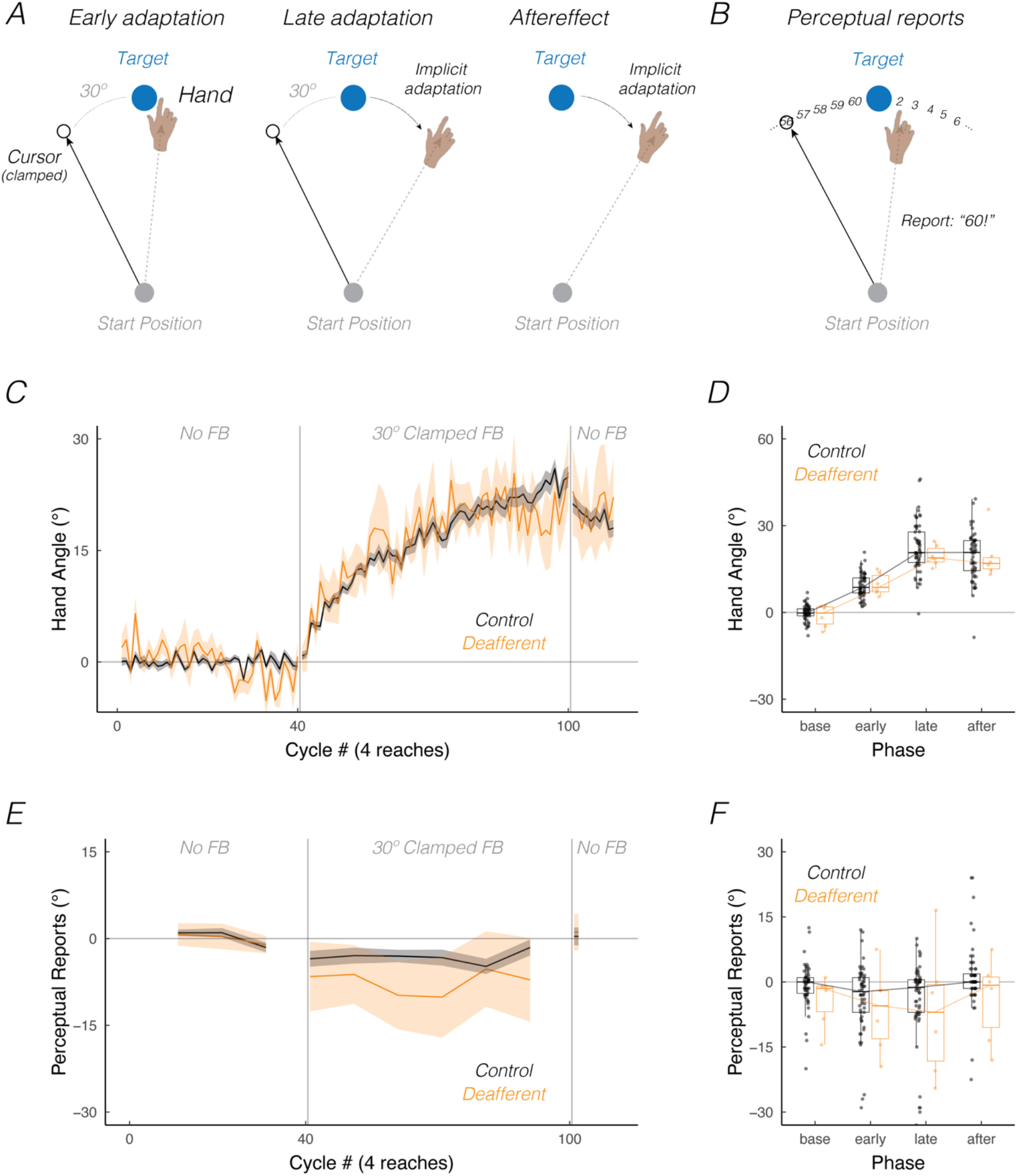
Minimal impact of proprioceptive loss on implicit motor adaptation and perceived movement outcome. **(A)** Schematic of the visual clamped feedback task. After baseline trials without cursor feedback (cycles 1 – 40), participants were exposed to 240 trials with clamped visual feedback (cycles 41 – 100) in which the cursor (white circle) followed a fixed trajectory, rotated 30° counterclockwise relative to the target. Participants were instructed to always move directly to the target (blue circle) and ignore the visual clamped feedback. Left, middle and right panels are schematics of hand and cursor positions during the early (cycles 41-60), late (cycles 81-100), and aftereffect (cycles 101-110) phases of adaptation, respectively. **(B)** Every 10^th^ cycle, participants reported their perceived movement outcome. On these trials, a number wheel would appear on the screen as soon as the amplitude of the movement reached the target distance, cueing participants for a report (top panel). The numbers (“1” to “60”) increased incrementally in the clockwise direction (spaced at 6° intervals around the circle), with the number “1” positioned at the target location. Participants used their keyboard to type the number closest to their perceived movement outcome when reaching the target distance. Mean time courses of hand angle **(C)** and perceptual reports **(E)** for Control (black; N = 60) and Deafferented groups (orange; N = 6). Shaded areas represent standard error. Both measures are presented relative to the target (0°); negative and positive values denote movements/reports toward or away from the cursor, respectively. One cycle consisted of four movements, one to each of the four possible target locations. Summary of implicit adaptation **(D)** and perceptual report data **(F)** over baseline, early, late and aftereffect phases. Box plots show minimum, median, maximum, and 1st/3rd interquartile values. Dots denote median for each individual.

Consistent with previous studies using the clamped feedback task, the control group showed a gradual change in hand angle in the opposite direction to the 30° clamped visual feedback, with the deviation averaging ∼20° away from the target at the end of the clamped feedback block (Figure 1C). The deafferented group showed a similar pattern of adaptation. These data provide a compelling demonstration that implicit adaptation is preserved despite the loss of proprioceptive and tactile afferents.

We analyzed the data at four phases in the experiment: baseline (with veridical visual feedback), early adaptation (with clamped feedback), late adaptation (with clamped feedback), and aftereffect (with no visual feedback). There was a main effect of phase (*F*(3, 192) = 287.0, *p* < 0.001, *η*^2^ = 0.61). Implicit adaptation was observed during the early, late, and aftereffect phases (all phases vs baseline hand angle, *t*(192) > 11.0, *p* < 0.001, *D*_*z*_ > 2.8) (Figure 1D). Hand angle increased from early to late adaptation (*t*(192) = 15.3, *p* < 0.001, *D*_*z*_ = 2.0). When visual feedback was eliminated during the aftereffect block, hand angle remained elevated, exhibiting only a small decrease compared to that observed late in adaptation (*t*(192) = −3.4, *p* < 0.001, *D*_*z*_ = 0.6). This result highlights that the change in hand angle elicited by clamped feedback was implicit.

Turning to our main question, we did not observe any significant differences in the extent of implicit adaptation between the deafferented and control participants. There was neither a significant main effect of group (*F*(1, 159) = 0.99, *p* = 0.32, *η*^2^ = 0.00, *BF*_01_ = 4.0, moderate evidence for the null) nor a significant interaction between group and phase (*F*(3, 192) = 0.5, *p* = 0.68, *η*^2^ = 0.00, *BF*_01_ = 6.2, moderate evidence for the null). Notably, *all* of the deafferented participants exhibited a substantial aftereffect, underscoring that implicit adaptation is preserved in this population. However, there was no evidence that proprioceptive loss heightened implicit adaptation.

### Perceptual shift is preserved but not heightened in deafferented individuals

We next turned to the question of how perceived movement outcome was impacted by proprioceptive loss. Every 10^th^ movement cycle, a number wheel appeared on the screen immediately after the center-out reaching movement was completed (Figure 1B). Similar to that of previous studies (Tsay et al., 2020), participants had to report their perceived movement outcome when the cursor crossed the target distance; to do this, they used the computer keyboard to type in the number closest to their perceived movement position. Following the report, the white cursor reappeared at a random position near the start position, cueing the participant to move the cursor back to the start position to initiate the next trial.

Perceptual reports were unbiased in baseline (denoted by near zero reports in Figure 1E) and exhibited a shift toward the perturbed visual feedback during the clamped feedback block (denoted by negative reports). This perceptual shift, present even after only one clamped feedback cycle, can be considered to result in a perceptual error given the assumption that the desired movement position is at the target (per task instructions). For the control participants, the perceptual error remained relatively constant across most of the adaptation block, only re-aligning back to the target at the end of the late adaptation phase. The deafferented group also showed a shift toward the perturbed visual feedback with the onset of the perturbed feedback, and this shift persisted throughout the adaptation block.

We analyzed the data at four phases in the experiment: baseline, early adaptation, late adaptation, and aftereffect phases. There was a main effect of phase (*F*(3, 192) = 7.3, *p* < 0.001, *η*^2^ = 0.03). Compared to the baseline phase, perceived movement outcomes in both groups were significantly (but subtly) biased toward the visual cursor during early and late adaptation phases (early vs baseline reports: *t*(192) = −2.4, *p* = 0.02, *D*_*z*_ = 0.3; late vs baseline reports: *t*(192) = −2.5, *p* = 0.01, *D*_*z*_ = 0.3). The Control group exhibited a -3.3 ± 1.1° perceptual shift (*p* = 0.02) (i.e., change in perceived movement outcome between early and baseline phases), a value consistent with prior work (Tsay et al., 2020). Notably, the Deafferented group shifted -7.5° ± 4.5° in the same direction (*p* = 0.04), with all but one (IW) deafferented participant exhibiting this perceptual shift (Figure 1F). The magnitude of the shift was similar in the two adaptation phases (early vs late: *t*(192) = −1.7, *p* = 0.87, *D*_*z*_ = 0.03), but dissipated when visual feedback was removed in the aftereffect phase (late vs aftereffect: *t*(192) = 4.1, *p* < 0.001, *D*_*z*_ = 0.6; aftereffect vs baseline: *t*(192) = 1.5, *p* = 0.13, *D*_*z*_ = 0.2).

Turning to the comparison between groups, we did not observe any significant differences in perceptual reports between the deafferented and control participants. There was neither a significant main effect of Group (*F*(1, 216) = 0.9, *p* = 0.34, *η*^2^ = 0.04, *BF*_01_ = 0.5; anecdotal evidence in favor of the null) nor a significant interaction between Group and Phase (*F*(3, 192) = 0.95, *p* = 0.95, *η*^2^ = 0.0, *BF*_01_ = 6.8; strong evidence in favor of the null). Thus, deafferented individuals exhibited similar biases in perceived outcome as the control participants, resulting in a perceptual error. However, also at odds with our prediction, this shift was not heightened by proprioceptive loss.

While our findings indicate that proprioceptive loss has minimal impact on perceived movement outcome, there are several limitations to these perceptual reports, many of which we have outlined in the Supplemental Section: Limitations with Reports of Perceived Movement Outcome.

### Motor control impairments in deafferented individuals

To evaluate motor performance in deafferented individuals in this task, we focused on the kinematic data from the baseline phase, prior to the introduction of the perturbed feedback. As shown in Figure 2, there were no significant group differences in movement time (Control: 102.0 ± 10.1 ms, Deafferented: 92.3 ± 12.3 ms; *t*(14) = 0.6, *p* = 0.60, *D* = 0.1). Moreover, neither group showed a significant bias in reach angle during the baseline block (baseline vs 0: Controls, *t*(59) = −0.4, *p* = 0.69, *D*_*z*_ = 0.1; Deafferented: *t*(5) = −0.8, *p* = 0.46, *D*_*z*_ = 0.1). However, hand angle variability was larger in the Deafferented group compared to the Control group (signed hand angle SD: Control: 6.4 ± 0.4°, Deafferented: 8.9 ± 0.9°; *t*(7) = 2.7, *p* = 0.03, *D* = 0.9; un-signed hand angle SD: Control: 4.1 ± 0.3°, Deafferented: 5.3 ± 0.4°; *t*(11) = 2.5, *p* = 0.03, *D* = 0.6), indicating that movements were less consistent when proprioception was impaired.

**Figure 2:**
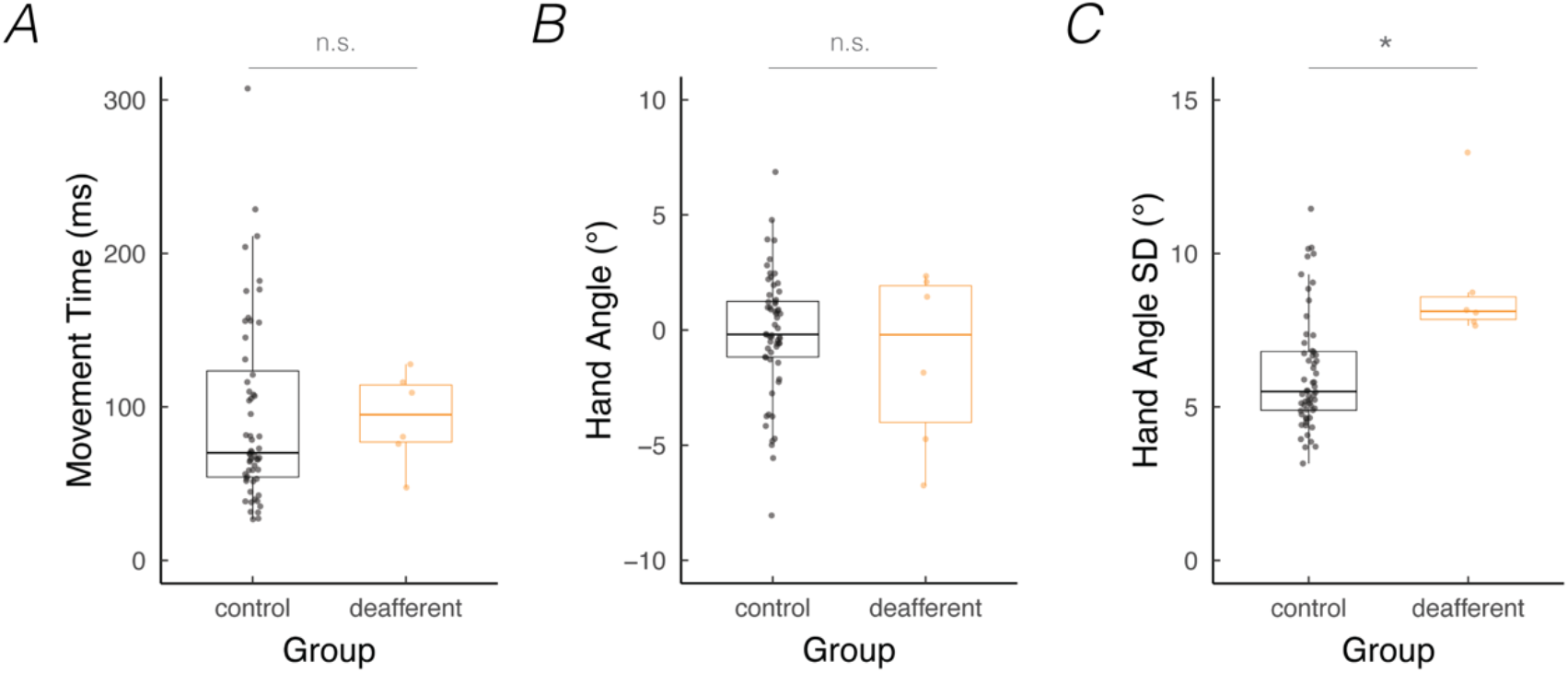
Proprioceptive loss results in greater motor variability. **(A)** Movement time, **(B)** mean hand angle, and **(C)** hand angle variability (i.e, standard deviation of unsigned hand angles) during baseline no-feedback trials in deafferented individuals (orange) compared their matched controls (black). Box plots show minimum, median, maximum, and 1st/3rd interquartile values. Dots denote individuals. ^*^ denotes p<0.05.

Given this difference, we repeated our between-group analysis of implicit adaptation and included hand angle variability as a covariate. There was neither a significant main effect of Group (*F*(1, 154) = 0.1, *p* = 0.74, *η*^2^ = 0.00), nor a significant interaction between Variability and Group (*F*(3, 193) = 0.7, *p* = 0.57, *η*^2^ = 0.01). Thus, the preservation of implicit adaptation and perceptual shift in deafferented adults was observed despite their increased variability.

## Discussion

Individuals lacking proprioceptive and tactile inputs provide an important test case for understanding the role of proprioception in implicit adaptation. While previous studies have observed preserved motor adaptation in deafferented adults (Bernier et al., 2006; Ingram et al., 2000; Lefumat et al., 2016; Miall et al., 2018; Sarlegna et al., 2010; Yousif et al., 2015), the motor tasks employed did not isolate implicit adaptation. To address this, we used a modified visuomotor rotation task to cleanly examine implicit motor adaptation and probe perceived movement outcome in deafferented adults. We found that the deafferented group exhibited robust implicit adaptation and perceptual shifts toward the visual perturbation. Moreover, we did not observe any differences on these measures between the deafferented and control groups. These findings underscore how proprioceptive loss has minimal impact on the extent of implicit motor adaptation and perceived movement outcome.

Our study is the first, to the best of our knowledge, to examine perceived movement outcome during motor adaptation in deafferented participants. We expect this question has not been asked because it may seem odd to probe changes in perceived movement position in participants who lack proprioception. Interestingly, none of our participants found making perceptual reports unintuitive or difficult. This underscores how proprioception may not be necessary for these perceptual reports, given that this multimodal percept also relies on visual and predictive signals (i.e., prior expectations from the intended aim and the efferent motor command) (Desmurget & Grafton, 2000; Gandevia et al., 2006; Proske & Gandevia, 2012; Wolpert et al., 1995). As such, we hypothesized that loss of proprioception would lead to a heightened dependence on vision when determining perceived outcome. However, the magnitude of perceptual shifts did not statistically differ between the control and deafferented groups.

We also hypothesized that proprioceptive loss would lead to heightened implicit adaptation in deafferented adults, to offset a heightened perceptual error. Given that the deafferented group did not show an increase in the perceptual shift, the current study does not provide a strong test of this prediction. Nonetheless, it is noteworthy that in the current study, as well as past work (Bernier et al., 2006; Ingram et al., 2000; Miall et al., 2018; Sarlegna et al., 2010; Sarlegna & Sainburg, 2009; Yousif et al., 2015), adaptation did not statistically differ between the control and deafferented groups.

The lack of significant differences between deafferented and control participants appears to align with a visuo-centric model of implicit adaptation. According to this view, implicit adaptation is driven by visual error – the difference between predicted and actual visual feedback (Burge et al., 2008; Morehead et al., 2017). Since this model does not include proprioception, deafferentation would not impact implicit adaptation. However, the model is not without shortcomings. It fails to explain why the magnitude of the perceptual shift correlates with the extent of implicit adaptation in prior studies (Salomonczyk et al., 2013; Tsay, Kim, et al., 2021).

Alternatively, implicit adaptation in deafferented adults might reflect the operation of compensatory mechanisms associated with chronic proprioceptive loss. As posited by the perceptual re-alignment model, proprioceptive afferent and efferent signals convey information about the hand position. With chronic proprioceptive loss, the perceived movement outcome might be primarily defined by the efferent motor command (Bard et al., 1999; Bernier et al., 2006; Fleury et al., 1995; Sarlegna et al., 2006). This post-hoc account of the null findings in the current study puts forth an important prediction: Assuming reweighting is a gradual process, a transient disruption in proprioception such as from muscle vibration (Goodwin et al., 1972) or non-invasive brain stimulation (Kumar et al., 2019; Ohashi et al., 2019) should enhance implicit adaptation.

## Methods

### Ethics Statement

All participants gave written informed consent in accordance with policies approved by the UC Berkeley’s Institutional Review Board. Participation in the study was in exchange for monetary compensation.

### Participants

We recruited deafferented participants who despite their severe upper-limb sensory loss, could perform a simple reaching task. Given the rarity of this combination, we used an online approach to test six chronic, deafferented participants spread across four countries (Tables 1-2). This sample is larger and more etiologically diverse than recruited in prior studies on this topic. While there are no gold standards for the clinical evaluation of proprioception, we obtained medical reports for each deafferented participant, all of which indicated that clinical assessments of proprioception were abnormal and upper-limb reflexes were impaired or absent.

**Table 1:** Deafferented participant demographics. Participants identified as either male (M) or female (F), right-handed (R) or left-handed (L).

| Name | Etiology | Age | Years since onset | Sex | Handedness |
| --- | --- | --- | --- | --- | --- |
| CD | Congenital | 22 | 22 | F | R |
| CM | Congenital | 46 | 46 | M | R |
| SB | Congenital | 34 | 34 | F | R |
| DC | Acquired | 54 | 16 | F | R |
| IW | Acquired | 70 | 51 | M | L |
| WL | Acquired | 53 | 22 | F | L |

**Table 2:**
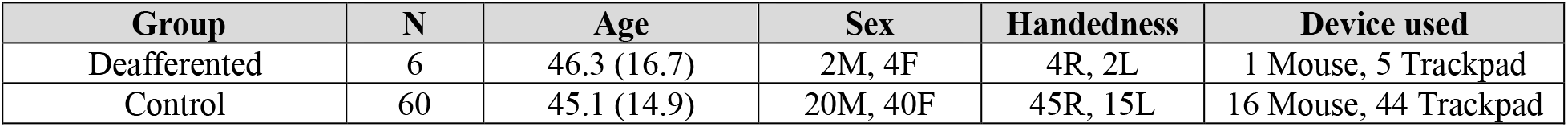
Deafferented and age, sex, handedness, and device-matched control participants. Participants either used a mouse or trackpad to complete the experiment. The two groups were well-matched on multiple dimensions (Age: *t*(6) = 0.2, *p* = 0.86, *D* = 0.1; Sex (M: male or F: female).: *χ*^2^(1, 66) = 0, *p* = 1; Handedness (R: right-handed or L: left-handed): *χ*^2^(1, 66) = 0.2, *p* = 0.66; Device used: *χ*^2^(1, 66) = 0.3, *p* = 0.59).

In terms of etiology, three participants have a congenital disorder that affects proprioception and tactile perception, and results in severe motor ataxia: CM and SB have an autosomal recessive mutation in the mechanoreceptor PIEZO2 gene (Chesler et al., 2016). CD has an inherited mutation in the mechanoreceptor ASIC3 gene (Lin et al., 2016). The three other participants had acquired deafferentation following an acute neurological episode. IW suffered a sensory neuropathy at age 19 from an autoimmune response to a viral infection, resulting in severe proprioceptive and tactile impairment below the neck (Cole & Katifi, 1991; Cole & Sedgwick, 1992). WL had a bout of polyradiculitis at age 31 which resulted in severe proprioceptive and tactile impairments below the neck (Miall et al., 2018, 2019). DC has severe proprioceptive impairment in the right upper limb subsequent to surgical resection at age 38 of a vascular tumor near the right medulla oblongata (Cardinali et al., 2016; Miller et al., 2019).

A total of 60 control participants were recruited, with 10 controls selected to match each of the deafferented participants in terms of age, sex, handedness, and device used in the experiment (Table 2). Control participants were recruited via Prolific, an online crowdsourcing platform connecting researchers to willing participants around the world.

The deafferented participants completed the task during a live video session, with the experimenter available to provide instructions and monitor performance. The control participants completed the task autonomously, accessing the website at their convenience.

### Apparatus

Participants used their own computer to access a dynamic webpage (HTML, JavaScript, and CSS) hosted on Google Firebase (Tsay, Lee, et al., 2021). The task progression was controlled by JavaScript code running locally in the participant’s web browser. The participant’s screen size was automatically detected, and this information was used to scale the size and position of the stimuli. There was no significant difference in screen size between groups (height: *t*(9) = 0.4, *p* = 0.71, *D* = 0.1; width: *t*(10) = 1.8, *p* = 0.10, *D* = 0.6). For ease of exposition, the parameters below are based on the average screen size (width x height: 1455 x 831 pixels).

We note that, unlike our laboratory-based setup in which we occlude vision of the reaching hand, this was not possible with the online testing protocol. That being said, we have found that measures of implicit adaptation are similar between in-person and online settings (Tsay, Lee, et al., 2021). Moreover, based on our informal observations, participants remain focused on the screen during the experiment (to see the target and how well they are doing) and did not appear to directly gaze at their hand.

### Procedure

Participants used either a trackpad or mouse to move a computer cursor (see a video describing the task here: https://youtu.be/6eJ78sQsjF8). Participants made a center-out movement from the center of the workspace to a visual target. A white annulus (0.5 cm in diameter) indicated the center position, a blue circle indicated the target location (0.5 cm in diameter), and the cursor was a white dot (0.5 cm in diameter). There were four possible target locations equally spaced around the workspace (45°, 135°, 225°, 315° where 0° corresponds to the rightward direction). On each trial, the target location was selected in a pseudo-randomized manner, with each target appearing once every cycle of four trials. The radial distance of the target from the start location was 8 cm on the visual display. The physical movement distance was likely between 6 cm – 10 cm (set to fit within the perimeter of the trackpad/tabletop), determined by the sensitivity (gain) setting of the participants’ device. Participants’ movements were limited to the wrist and fingers given the device used and required movement distance. Prior to starting the experiment, participants had to watch an instructional video, which provided an overview of the procedure.

At the beginning of each trial, the cursor appeared at a random position within 1 cm of the center of the screen. As such, the actual starting hand position varied subtly from trial to trial. The participant initiated the trial by moving the cursor to the center start location. After maintaining the cursor in the start position for 500 ms, the target appeared. Participants were instructed to move rapidly, attempting to “slice” through the target. There were three types of feedback conditions during the experiment: No visual feedback, veridical visual feedback, and clamped visual feedback. During no-feedback trials, the cursor was extinguished as soon as the hand left the start annulus and remained off for the entire reach. During veridical feedback trials, the movement direction of the cursor was veridical with respect to the movement direction of the hand. The veridical cursor was extinguished when the hand crossed the radial target distance of 8 centimeters. Note that veridical feedback trials were only used at the beginning of the experiment to familiarize the participant with the task. During clamped feedback trials (Figure 1A), the cursor moved at a 30° angular offset relative to the position of the target, counterclockwise and irrespective of the actual movement direction of the hand – a manipulation shown to isolate implicit adaptation (Morehead et al., 2017; Tsay et al., 2020). The clamped cursor was extinguished when the hand crossed the radial target distance of 8 centimeters.

Every 10th cycle, participants were asked to report their perceived movement outcome for four consecutive trials (i.e., one report per target location). There was a total of 40 ‘perceptual report’ trials over the course of the experiment. On perceptual report trials, a number wheel appeared on the screen as soon as the clamped cursor reached the target amplitude, cueing the participant for a report. The numbers (“1” to “60”) increased incrementally in the clockwise direction (spaced at 6° intervals around the circle), with the number “1” positioned at the target location. The participant used the keyboard to report the number closest to their perceived movement position. Following the report, the white cursor appeared at a random position within 1 cm of the center start position. The participant moved the cursor to the start position to initiate the next trial.

The main task consisted of 110 cycles (four reaches per cycle, 440 trials total) distributed across three main blocks of cycles/trials: A no-feedback block (40 cycles; 160 trials to assess baseline performance), clamped feedback block (60 cycles; 240 trials to assess adaptation), and a no-feedback block (10 cycles; 40 trials to assess aftereffects). Prior to the clamped feedback block, the following instructions were provided: “The white cursor will no longer be under your control. Please ignore the white cursor and continue to aim directly towards the target.”

To clarify the invariant nature of the clamped feedback, eight demonstration trials were provided before the first perturbation block. On all eight trials, the target either appeared straight ahead (90° position), and the participant was told to reach to the left, to the right, and backward. On all of these demonstration trials, the cursor moved in a straight line, 90° offset from the target. In this way, the participant could see that the spatial trajectory of the cursor was unrelated to their own reach direction.

To verify that the participants understood the clamped visual feedback manipulation task, we included an instruction check after eight demonstration trials in the adaptation block. The following sentence was presented on the screen: “Identify the correct statement. Press ‘a’: I will aim away from the target and ignore the white dot. Press ‘b’: I will aim directly towards the target location and ignore the white dot.” The experiment only progressed if participants pressed the “b” key.

### Data analysis

The main dependent variable for measuring adaptation was hand angle, defined as the angle of the hand relative to the target when movement amplitude reached 8 cm from the start position. This measure defines the angular difference between the target location and movement direction. Pilot work using our web-based platform indicated that reaching trajectories are generally fast and straight without evidence of online feedback corrections.

We defined four phases of adaptation: Baseline, early adaptation, late adaptation, and aftereffect. Baseline performance was operationalized as the mean hand angle over the no-feedback baseline block (cycles 1 – 40). Early adaptation was operationalized as the mean hand angle over the first 20 cycles of the clamped visual feedback block (cycles 41 – 60). Late adaptation was defined as the mean hand angle over the last 20 cycles of the clamped visual feedback block (cycles 81 – 100). The aftereffect was operationalized as the mean hand angle over the 10 cycles of the no-feedback aftereffect block (cycles 101 – 110).

Outlier responses were defined as trials in which the hand angle was greater than 90° from the target or deviated more than three standard deviations from a trendline constructed with a moving 5-trial window. Outlier trials were excluded from further analysis since behavior on these trials could reflect anticipatory movements to the wrong target location or attentional lapses (average excluded movement trials: Control group = 1.3 ± 0.2%; Deafferented group = 1.1 ± 0.3%).

The perceptual reports provide the dependent variable for measuring the perceived movement outcome. These data were converted into angular values, although we note that the perceptual reports involve categorical data (numbers spaced at 6° intervals), whereas in angular form they suggest a continuous variable. Outlier responses were removed in the exact same manner as the hand angle data (average excluded report trials: Control group = 1.8 ± 1.0%; Deafferented group = 0.4 ± 0.4%). Variability in the perceptual reports did not differ between Control and Deafferented groups (Mean ± SEM, Control: 14.2° ± 2.2°; Deafferent: 15.7° ± 2.8°; *t*(13) = 0.4, *p* = 0.67, *D* = 0.1).

Reaction time was defined as the time from target presentation to the start of movement, defined as when the radial movement of the hand exceeded 1 cm of movement. Movement time was defined as the time between the start of movement and when the radial extent of the visual cursor (either hidden or provided) reached 8 cm, the target distance. If the movement time exceeded 500 ms, the message, “too slow” was displayed at the center of the screen for 750 ms before the next trial began.

Data were statistically analyzed using a linear mixed effect model (R: lmer function) with Phase (baseline, early, late, and aftereffect) and Group (Control and Deafferented) as fixed (interacting) factors and Participant as a random factor. Post hoc two-tailed t-tests on the betas from the linear mixed effect model were evaluated using the *emmeans* and *ANOVA* functions in R (Bonferroni corrected for multiple comparisons). Given the differences in sample size and group characteristics, we opted to use Welch’s t-tests. This test is designed for comparing two independent groups when it cannot be assumed that the two groups have equal variances. Standard effect sizes are reported (*η*^2^ for fixed factors; Cohen’s *D*_*z*_ for within-subjects t-tests, Cohen’s *D* for between-subjects t-tests).

### Supplemental Section: Limitations with Reports of Perceived Movement Outcome

In previous studies involving reports of perceived movement outcome during adaptation, the perceptual shift reached a maximum value shortly after the onset of the visual perturbation and then dissipated over time, returning to baseline levels in the last phase of adaptation (Synofzik et al., 2006; Tsay et al., 2020). Although this pattern was evident in the mean data for the control participants, there was no statistical reduction in the perceptual error (shift) between the early and late adaptation phases. Several factors might account for this observation: First, the study’s duration may have been too short. Specifically, our experiment spanned 1.5 hours and consisted of 440 trials. This design choice was made to minimize fatigue and cater to the mobility challenges faced by the deafferented participants. Extending the number of learning trials and perceptual probes may clarify whether the perceptual error diminishes as implicit adaptation ceases.

Second, the perceptual probes in our study may have been subject to unaccounted influences such as gaze direction (Jones & Henriques, 2010), transformations across horizontal and vertical workspaces, participants’ interpretations of the directive to “ignore the visual cursor”, and the presence of a visual target. That is, participants, both controls and patients, might have been inclined to base their perceptual reports on the location of the visual target and clamped visual feedback, rather than on efferent and/or proprioceptive feedback conveying hand position. To obtain a more precise measure, future studies could examine perceived movement outcome after a passive or self-initiated movement, in the absence of both a visual cursor and target (Izawa & Shadmehr, 2011).

We also failed to find heightened perceptual shifts in deafferented adults. There are at least two potential explanations for this null result. First, although the deafferentation was severe, the proximal muscles of the upper extremity were possibly spared in a subset of our participants. As such, residual proprioceptive input might have limited the magnitude of perceptual shifts toward vision. This possibility seems unlikely given that our task predominantly involved distal finger/wrist movements. While cognizant of the small sample size, we did not observe any relationship between the clinical evaluation and size of the perceptual shift.

Second, while the online format of our study enabled the recruitment of a sizable deafferented cohort, it did require some changes to the standard methods used to test implicit motor adaptation and perceived movement outcome. The perceptual judgments were made following an active movement, rather than a passive movement of the hand (via a robot or experimenter). Furthermore, participants had peripheral vision of their actual hand position, and such visual input could impact both adaptation and the perceptual reports. Although our data on control participants suggest that implicit adaptation occurs whether it is measured online or in person (Tsay, Lee, et al., 2021), future experiments should examine the issues of visual feedback and passive movements.

## Footnotes

* In the original exposition of this model, we used the term “proprioceptive re-alignment”. However, recognizing that perceived movement outcome is influenced by feedback from vision and proprioception, the prior expectation conveying the intended action, and the efferent motor command, (Desmurget & Grafton, 2000; Gandevia et al., 2006; Proske & Gandevia, 2012; Wolpert et al., 1995) – a point made salient by Zhang et al (Zhang et al., 2023) – we now adopt the phrase “perceptual re-alignment” to better capture this idea.

